# A machine learning approach to predict drug-induced autoimmunity using transcriptional data

**DOI:** 10.1101/2023.04.04.533417

**Authors:** Gemma L. Smith, Ieuan G. Walker, Anna Aubareda, Michael A. Chapman

## Abstract

Drug-induced autoimmunity (DIA) is an idiosyncratic adverse drug reaction. Although first reported in the mid-1940’s, the mechanisms underlying DIA remain unclear, and there is little understanding of why it is only associated with some drugs. Because it only occurs in a small number of patients, DIA is not normally detected until a drug has reached the market. We describe an ensemble machine learning approach using transcriptional data to predict DIA. The genes comprising the signature implicate dysregulation of cell cycling or proliferation as part of the mechanism of DIA. This approach could be adapted by pharmaceutical companies as an additional preclinical safety screen, reducing the risk of drugs with the potential to cause autoimmunity reaching the market.

## Introduction

### Challenges of drug-induced autoimmunity

Autoimmune diseases arise when the immune system attacks the cells of an individual in error, due to a loss of tolerance of self-antigens. Autoimmune diseases can either target a specific organ, as seen in Type 1 diabetes, or cause systemic illness, as seen in systemic lupus erythematosus, and may be auto-antibody or cell mediated (Amital and Shoenfeld, 2004). Autoimmune disease occurs when a genetically predisposed individual is exposed to an environmental trigger. Triggers include viral, bacterial, and parasitic infections, food components, vaccination, occupational and other chemical exposures, and pharmaceutical drugs (Molina and Shoenfeld, 2005). When a drug is confirmed to be the environmental trigger of autoimmunity, it is referred to as drug-induced autoimmunity (DIA).

DIA is an idiosyncratic adverse drug reaction (ADR) in which administration of a drug causes one or more autoimmune symptoms, which are resolved following removal of the drug. Drugs from various classes have been shown to cause autoimmunity, and the range of autoimmune symptoms reported is large, including drug-induced lupus, autoimmune hepatitis, haemolytic anaemia, rheumatoid arthritis, membranous glomerulonephritis, myasthenia gravis, polymyositis, thyroiditis, Sjogren’s syndrome, scleroderma, dermatomyositis, pemphigus and pemphigoid (Xiao and Chang, 2014). A further complication of DIA is that a single drug can cause multiple autoimmune symptoms, as is seen with simvastatin which has been associated with lupus, dermatomyositis, polymyositis, and pemphigoid (Noël, 2007; Stoebner *et al*., 2003). DIA has a significant and widespread impact on society, causing approximately 10 % of all new lupus cases reported in the United States every year (Vedove *et al*., 2009). Additionally, patients being treated for relatively mild conditions can experience severe disease, as shown in the case of minocycline, an antibiotic prescribed to treat acne, which has been reported to induce lupus-like syndrome (Schlienger, Bircher and Meier, 2000). DIA can also cause substantial economic impact, as it is not normally detected until a drug has reached the market, and the severity of the diseases caused can lead to market withdrawal (Pichler, 2003). DIA occurs in a dose-independent manner, may take several months to manifest, can present in a similar manner to the idiopathic form of disease, and only occurs in a very small number of genetically predisposed patients (Dedeoglu, 2009). These factors not only make it difficult to diagnose DIA, but also to determine a potential mechanism of action and predict whether a drug is likely to cause DIA. Despite first being reported in 1945, when sulfadiazine was reported to cause lupus erythematosus (Hoffman, 1945), and over 100 drugs now being associated with DIA, no clear mechanism has been defined.

### Previous attempts to predict DIA

Many methods aiming to predict adverse drug reactions rely on the chemical structure of drugs, and this is a major focus of many attempts to predict DIA. Machine learning models using information on the chemical structure of drugs and reactive metabolite formation have been proposed to predict DIA. One such model demonstrated that drugs containing benzene with a nitrogen containing substituent were significantly associated with a higher risk of inducing autoimmunity (Wu *et al*., 2021). Others have compared different machine learning methods and different molecular fingerprint packages to predict DIA using molecular properties and structural alerts (Guo *et al*., 2022). Although helpful, chemical structure alone may not be sufficient to determine the risk of DIA in patients, and whilst structural analysis of chemical compounds is highly sensitive, it has the potential for generating false positives (Stepan *et al*., 2011).

### Use of gene transcription data to predict other adverse events

Advances in sequencing technology mean that obtaining gene expression data is now more affordable, and less time-consuming, resulting in many groups attempting to use gene expression data to predict adverse drug reactions. A machine learning classifier which combines both gene expression and the chemical structure of drugs has been used to predict adverse drug reactions (Wang, Clark and Ma’ayan, 2016). The LINCS L1000 dataset which contains gene expression data from various cell lines treated with multiple doses of a drug, and sampled at numerous timepoints, was used to generate this model. The drug/cell/dose/timepoint combination which gave the strongest gene expression signature compared to the control for each drug was used, as these datasets were the most predictive of ADRs. Genetic signatures were combined with information on chemical structure to develop a machine learning classifier capable of predicting ADRs, showing the value of such large datasets. They also demonstrated that gene profiling data was the best predictor of drug-ADR associations, compared to predictions from chemical structure or changes in cell morphology. Another attempt at using gene expression data to predict ADRs used the Side Effect Resource (SIDER) and the Connectivity Map (CMap) data to build a predictive “process-drug-side effect” network, which enables prediction of the cause of a side-effect based on the drugs known biological processes (Lee *et al*., 2011).

### Using transcriptional data to predict DIA

The SIDER database contains data on 1430 drugs and 5880 ADRs associated with those drugs (Kuhn *et al*., 2016). The Connectivity Map (CMap) is a database created by the Broad Institute that contains the gene expression signatures of hundreds of cell lines treated with ∼5,000 compounds (Lamb, 2006; Subramanian *et al*., 2017). Cell lines were treated with multiple concentrations of drugs, samples were taken at different timepoints, and their expression signature determined relative to a control. We proposed to explore the possibility of using these valuable datasets to find a gene expression signature predictive of DIA. However, we also foresaw potential challenges to this approach. For example, one would expect alterations in gene expression within the on-target pathway as well as in any off-target pathway responsible for initiation of autoimmunity. Furthermore, there are over 200 human cell lines from 21 different cell lineages in the CMap dataset and it was not clear that they would all contain the genetic machinery to trigger that off-target pathway. It was also not clear whether different drugs would target different off-target pathways, or whether different autoimmune conditions were triggered by different pathways. Finally, given that DIA often appears in a patient after weeks or months of treatment, it was not apparent whether we would see relevant changes during the 6- or 24-hour exposure times in CMap.

Perhaps surprisingly, we were able to detect positive signals associated with drugs that caused DIA, which we propose may be related to the ability of ensemble machine learning methods to capture complex patterns. But, despite this complexity, exploring our model revealed a common theme of disruption of genes involved in cell cycle regulation and proliferation, which we propose may relate to lowering the threshold for lymphocyte activation.

## Results

### Dataset

We downloaded the SIDER database and selected drugs with reports of autoimmunity as an adverse event registered with at least two registries. We manually curated a set of class-matched drugs that did not have reports of autoimmune side-effects, as a control. Because both statins and angiotensin-converting enzyme inhibitors (ACEIs) appeared to have autoimmunity as a class-wide side-effect, and it was not possible to select class-matched control drugs, we initially excluded these from the dataset. We then limited drugs to those that feature in CMap. This left 47 drugs with the potential to cause autoimmunity and 22 control drugs (Table 1). We then identified all perturbations in CMap that involved these drugs (a single perturbation is a combination of drug, dose, cell line, and exposure time). There were 1741 autoimmune drug perturbations and 1341 control drug perturbations (Fig. 1).

**Table 1.**
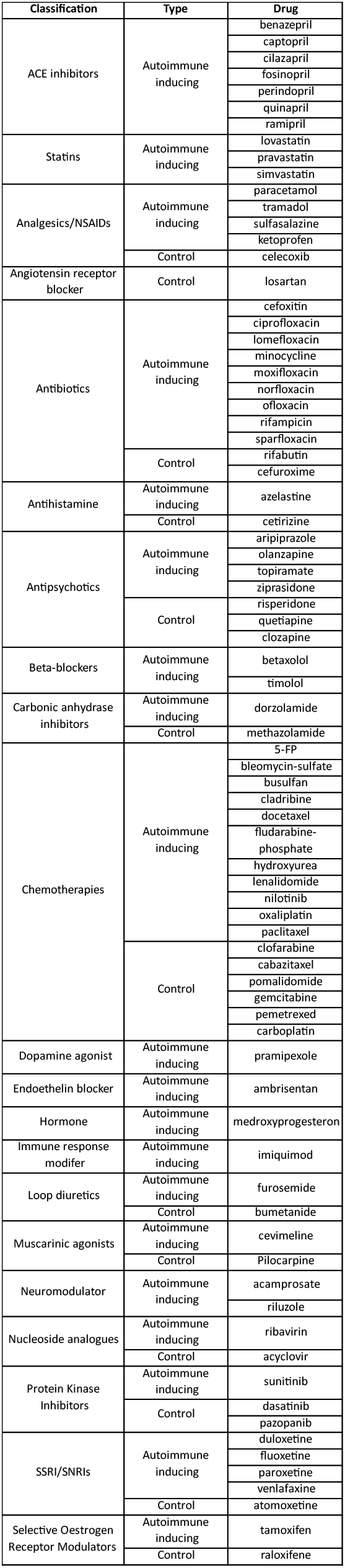
Details of the drugs used to train and validate the model. Note that statins and ACEIs were not included in the initial training dataset, as we could not identify a class-matched control.

**Figure 1.**
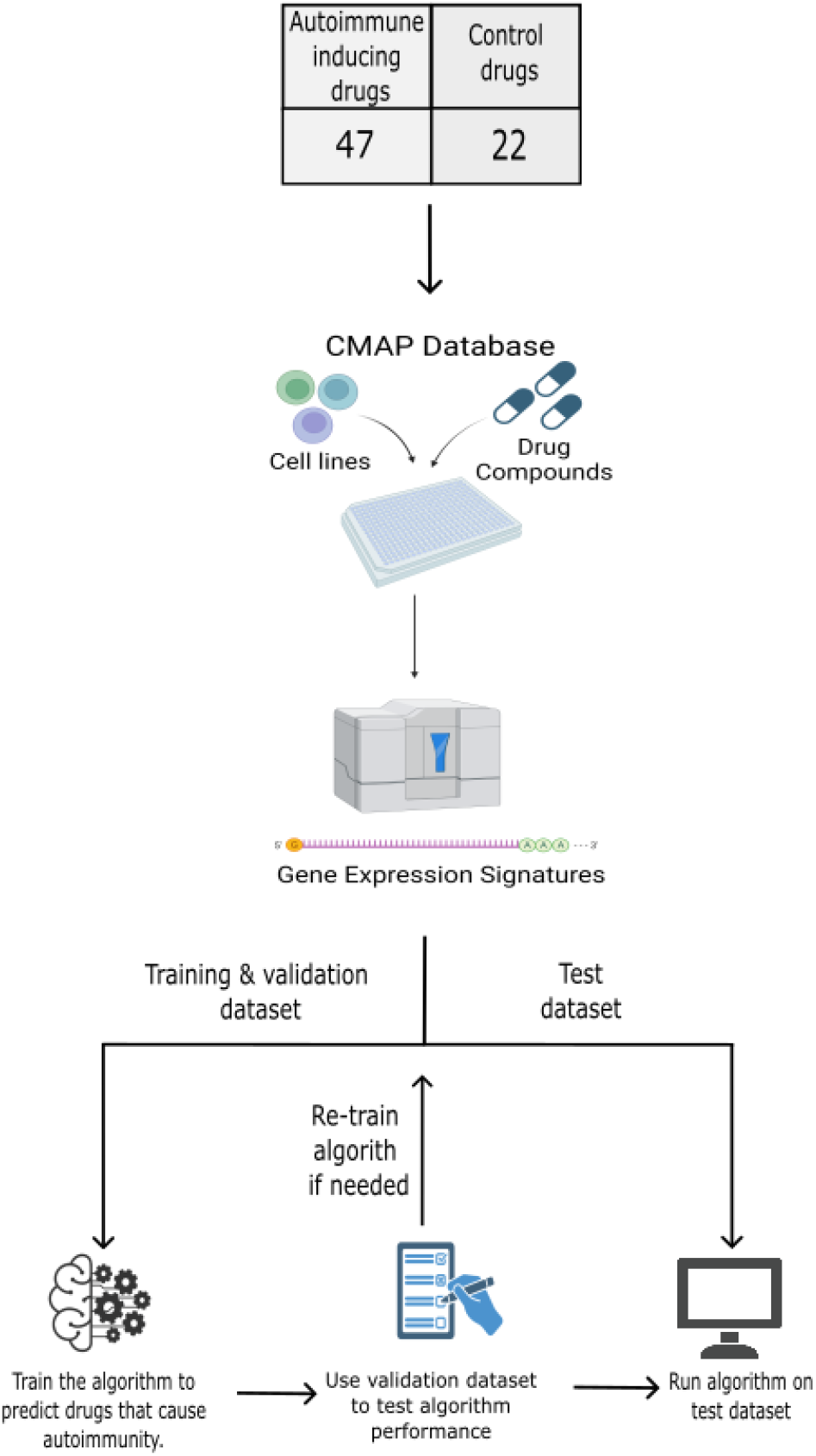
Overview of the process used to select data from the CMap database and train our machine learning model. The SIDER database was used to identify drugs with reports of autoimmunity as an adverse event registered with at least two registries, and a set of class-matched control drugs was manually curated. Statins and ACEIs were initially excluded from our analysis as a control could not be identified. We then limited drugs to those that feature in the CMap database, leaving 47 autoimmune inducing and 22 control drugs. We then extracted all perturbations for those drugs from the CMap, and divided them into training, validation, and test datasets, to be used in our machine learning algorithm.

### Potential for DIA cannot be captured by gene expression clustering structure

Different drug classes may precipitate the same autoimmune condition, and one drug class may induce multiple autoimmune conditions. However, it is not clear how many mechanisms are responsible for DIA across these different settings (Uetrecht, 2005). Furthermore, whilst any drug effect must result from a molecular alteration, and gene expression studies suggest that most molecular perturbations result in reproducible changes in gene expression patterns (Lamb, 2006), it is likely that such perturbation signatures will be cell lineage dependent. We therefore anticipated that when we clustered the 3082 CMap instances by gene expression, they would cluster into several sample clusters, reflecting this multiplicity of mechanisms. We then planned to explore these individual clusters for clues as to different mechanisms.

We adopted consensus clustering, a meta-clustering approach from which one can infer the number of stable underlying clusters (Monti *et al*., 2003). Expecting to see several stable clusters, we were surprised that consensus clustering implied a very stable two-cluster solution (Fig. 2A). Even when we specified up to five clusters, additional clusters were formed from a very few CMap instances, implying a lack of any real structure beyond the two-cluster solution (Fig. S1). It would have been highly unexpected had the two major clusters reflected autoimmunity potential, and indeed, this was not the case; instances involving drugs known to precipitate autoimmunity were not associated with either cluster (Fig. 2B; p=0.652, χ^2^-test). We also did not see any significant association between cell line and cluster membership (data not shown). Taken together, the clustering results confirm that simple differential gene expression approaches are unlikely to be helpful for identifying autoimmune-causing drugs from transcriptional perturbation data.

**Figure 2.**
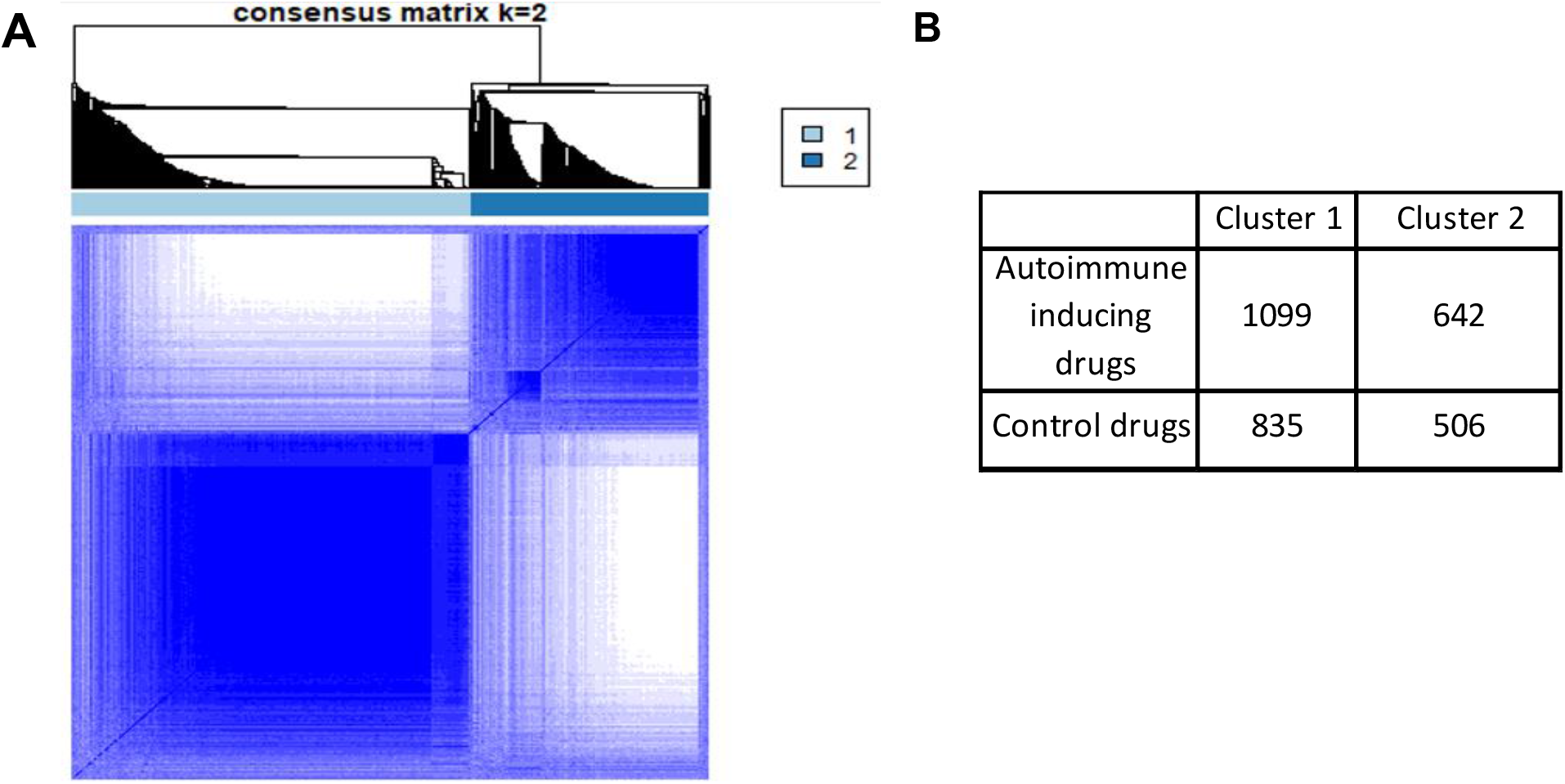
Consensus clustering of CMap instances does not separate drugs associated with autoimmunity from control drugs. A) Consensus clustering implies a very stable two-cluster solution for the selected CMap instances. B) A contingency table showing how the autoimmune-inducing and control drugs are distributed between the clusters.

### A regularised gradient boosting algorithm accurately classifies CMap instances by autoimmunity potential

We reasoned that a non-linear, ensemble machine learning approach might overcome issues of a multiplicity of drug effects and target cell line responses. XGBoost is a regularised gradient boosting algorithm that has proved highly successful in machine learning competitions (Chen and Guestrin, 2016). We divided the CMap instances of our autoimmune inducing drugs into 1000 learning instances, 250 validation instances, and 491 test instances. For the control drugs, there were 1000, 250, and 91 training, validation, and test CMap instances, respectively. We optimized XGBoost using the training and validation datasets, then predicted autoimmune potential of the CMap instances in the test dataset. The model demonstrated good discrimination ability with an area under the curve (AUC) of the receiver-operator characteristic (ROC) of 0.766 (Fig. 3A). We reasoned that some cell lines would contribute little to the discriminant ability of the model. When we confined testing to only those cell lines that had allowed accurate drug classification during validation, we saw an improvement during testing, with an AUC of 0.875 (Fig. 3B).

**Figure 3.**
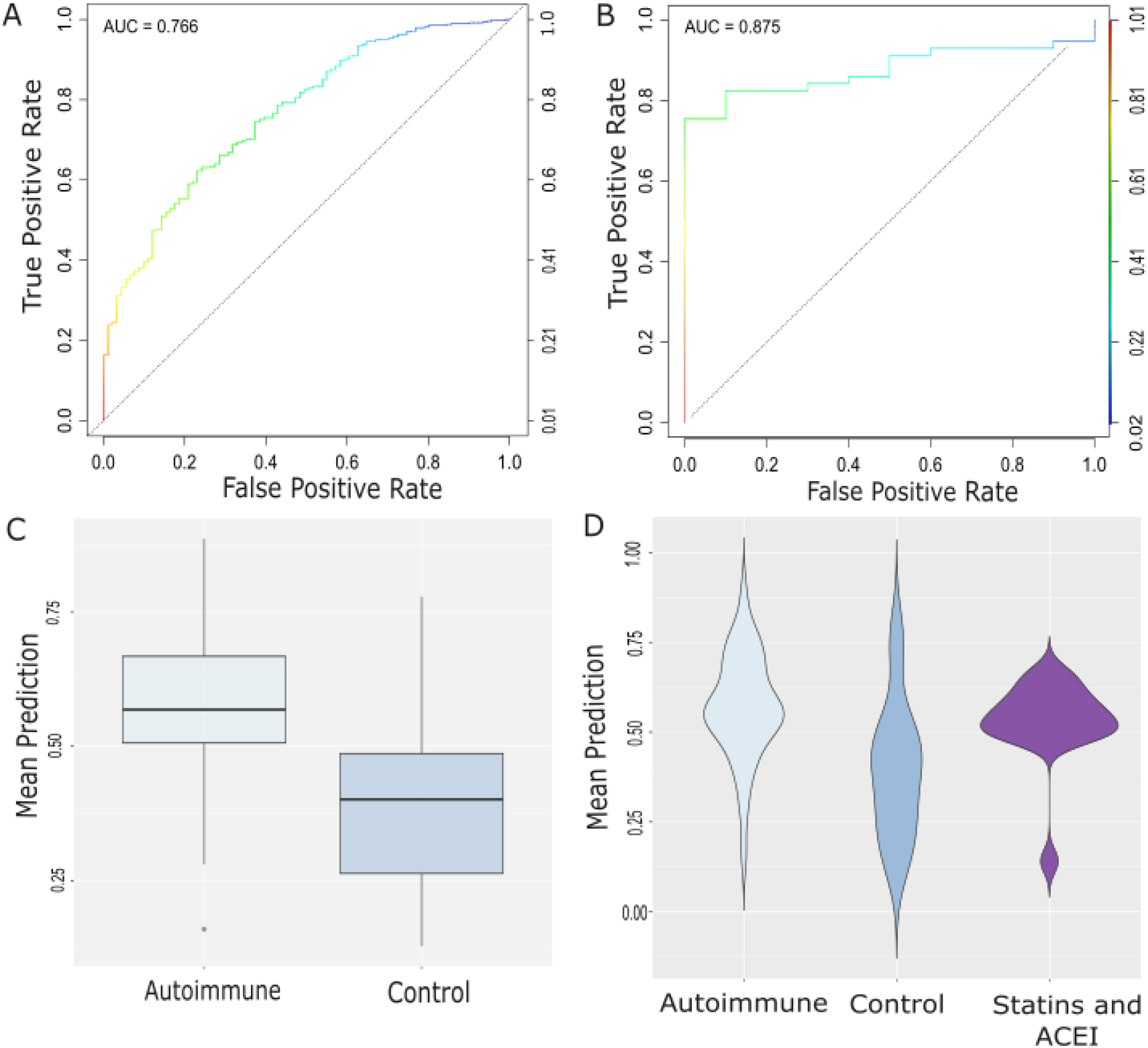
The optimised XGBoost algorithm accurately classifies CMap instances and drugs by autoimmune potential. A) Receiver-operator characteristic (ROC) curve for predicting CMap instances whose drugs were associated with autoimmunity. AUC: Area under the curve. B) ROC curve for predicting CMap instances whose drugs were associated with autoimmunity, considering only cell lines that performed well in validation. C) Per-drug scores for drugs associated with autoimmunity and control drugs. Scores are significantly different (p=0.003). D) Violin plots of per-drug scores as in (C) plus the per-drug scores of statins and ACEIs. Statins and ACEIs score significantly differently than control drugs (p=0.019) but not other autoimmune-inducing drugs (p=0.29).

In this last analysis, we were testing the ability of our model to classify individual instances, with multiple instances per drug. To make a per-compound classification, we reasoned that the predictions across all instances could be averaged out for each small molecule. To account for the varying reliability of the different cell lines noted above, one could take a weighted average of the prediction score, with weights provided by cell line accuracy during validation. We explored this approach, making a per-drug prediction for our test and control drugs. The per-drug scores were well-separated between our autoimmune inducing and control drug groups (Fig. 3C; p=0.003), and classification of the drugs achieved a sensitivity of 0.769 and a specificity of 0.812. Taken together, these results demonstrate that it is possible to use transcriptomic data to identify small molecules with the potential to induce autoimmune complications.

### The algorithm can distinguish previously un-encountered drug classes with autoimmunity potential

To prevent overfitting, we had ensured that the CMap instances constituting our test samples were independent from those in the training or validation sets. However, because there are multiple CMap instances per drug, there was overlap between drugs in the training, validation, and test datasets. We had omitted statins and ACEIs from the autoimmune inducing drug list because, in both cases, autoimmunity appears to be a class-wide effect and we were unable to find any class-matched control drugs. This also provided us the opportunity to test the ability of our model to classify drug classes to which it was completely naïve. At the per-drug level, the model correctly classified four out of seven ACEIs and seven out of nine statins as having the potential to cause autoimmunity. However, the output from the model is a continuous score, and even those statins and ACEIs not classified as autoimmune-causing scored close to the cut-off (with the exception of trandolopril). As a whole, the distribution of statin/ACEI classification scores were clearly shifted away from that of the control drugs (p=0.019; Fig. 3D), and was not significantly different to that of the autoimmune drugs (p=0.29; Fig. 3D). These data suggest that our model retains predictive power in drug classes to which it is naïve.

### The algorithm can distinguish drugs across multiple classes with the potential to trigger a previously un-encountered autoimmune condition

As a further test of the predictive ability of our model, we wanted to see how our model would perform classifying several drugs across multiple classes known to precipitate an autoimmune condition that had not been considered during training of the algorithm. Cutaneous lupus erythematosus (CLE) and systemic lupus erythematosus (SLE) are both autoimmune conditions that are commonly induced by drugs, relative to other autoimmune symptoms (He and Sawalha, 2018). However, we omitted lupus from our list of autoimmune conditions when selecting our drugs for training, to see if the algorithm could identify lupus-precipitating drugs, despite being naïve to these.

We first investigated how the model would score the majority of EMA/FDA-approved drugs. We took all CMap instances, where available, for all drugs in the SIDER database. There were 696 such drugs associated with 27159 CMap instances. The per-drug scores for these drugs ranged from 0.071 to 0.959 (Fig. 4A). We then compared the prediction scores of drugs with at least two separate registry reports of CLE or SLE in the SIDER database with all other drugs. This demonstrated that the prediction scores of lupus-precipitating drugs were significantly higher than those that had not been associated (p=0.002; Fig. 4B). These data further support the notion that our model can detect small molecules that are able to precipitate autoimmunity, even in previously unseen drug classes.

**Figure 4.**
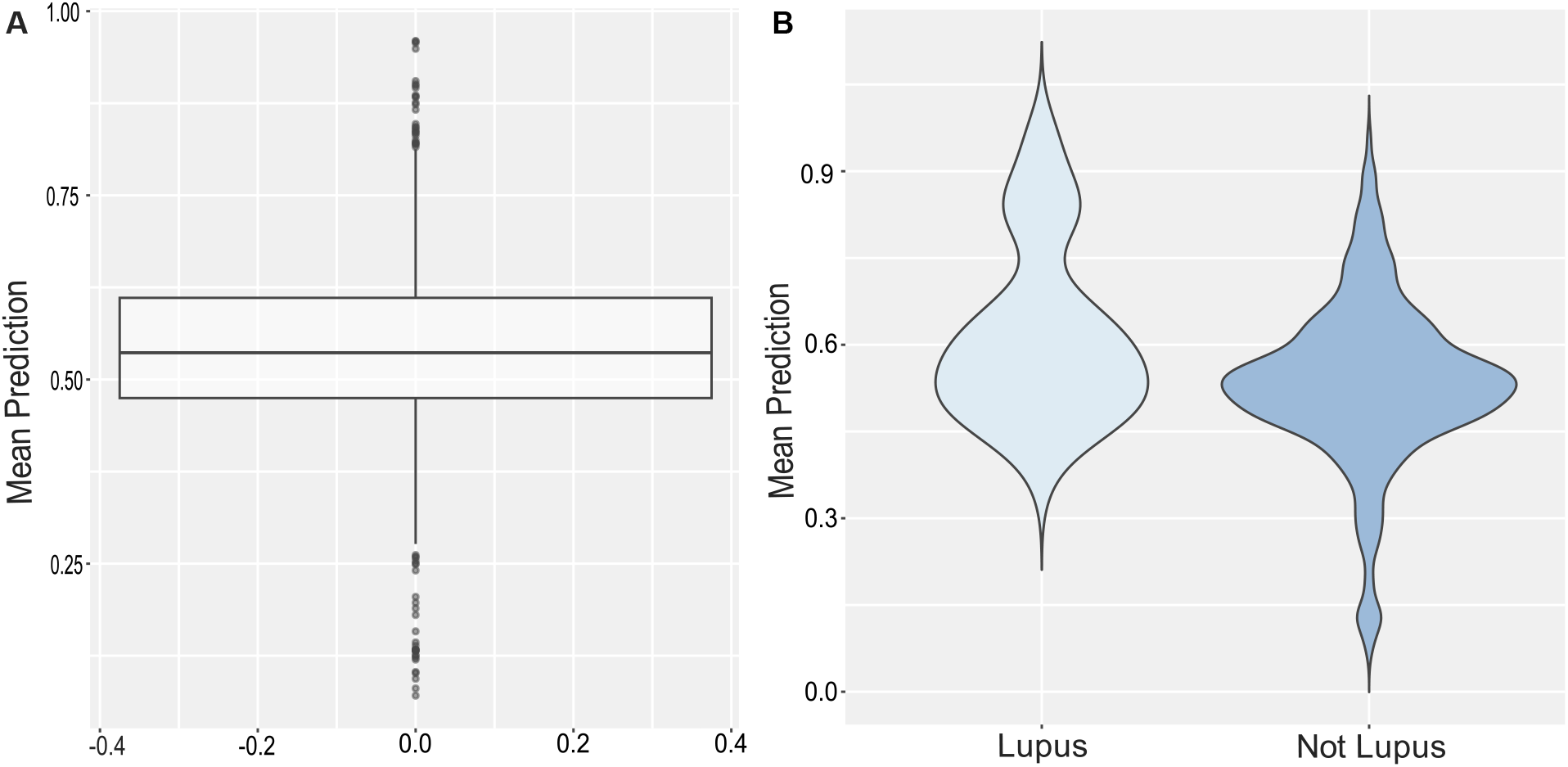
Drugs reported to cause SLE and CLE score higher than drugs not associated with lupus. A) Boxplot displaying the prediction scores given to all FDA/EMA approved drugs by the model. B) Violin plot of the prediction scores for drugs with at least two reports of SLE or CLE and for drugs not associated with lupus. The prediction score of drugs associated with lupus is significantly higher than other drugs (p = 0.002).

### A random forest machine learning approach has similar performance to regularised gradient boosting

Although the power of the XGBoost algorithm is well-documented, there is a risk that if a significant finding is only seen with one machine learning algorithm, that finding might be an artifact. We therefore repeated the analysis above with a related ensemble machine learning algorithm, the random forest (Breiman, 2001). This also relies on examining multiple trees, although the way it combines the information from each tree is different to XGBoost. The results of the analysis are shown in Fig. 5. As above, testing on CMap instances resulted in good classification, with an AUC of 0.779 (Fig. 5A), although confining testing to reliable cell lines did not noticeably improve classification, with an AUC of 0.794 (Fig. 5B). Per-drug scores were again well-separated (Fig. 5C; p=0.009), although per-drug sensitivity was lower than with XGBoost, at 0.650. Specificity was similar, at 0.765. Statins and ACEIs scored higher than control drugs (Fig. 5D; p=0.048), but did not have scores significantly different to autoimmune-precipitating drugs (Fig. 5D; p=0.511). When tested across a large number of approved drugs, the range of scores was lower, at 0.293-0.729 (Fig. 5E), than with our XGBoost approach. Again, drugs associated with CLE or SLE scored higher than those not associated with lupus (Fig. 5F; p=0.010). Taken together, these data demonstrate that a different tree-based machine learning algorithm also has predictive power for the potential of small molecules to precipitate autoimmunity and that this feature is not an isolated artifact of XGBoost.

**Figure 5.**
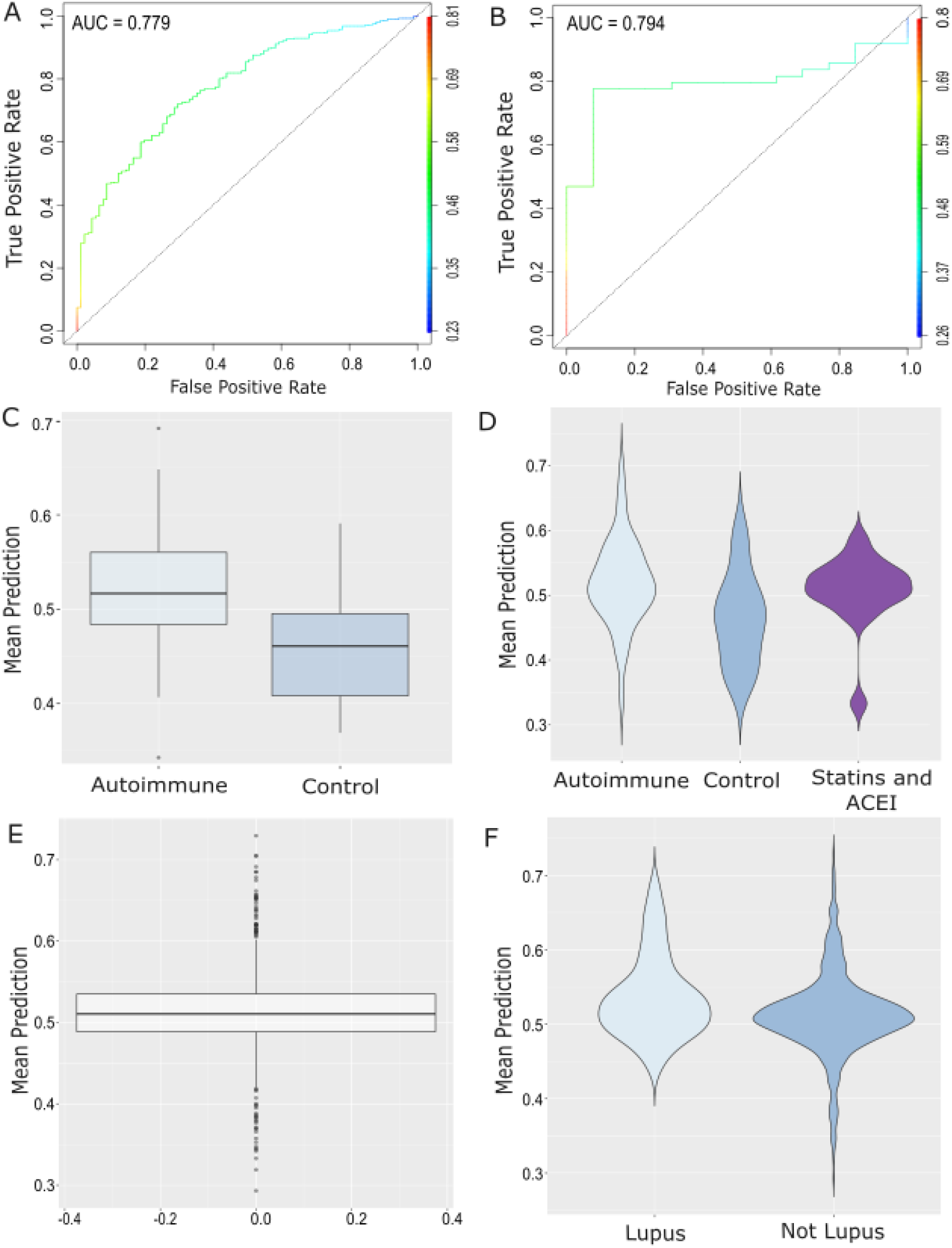
A random forest model performs similarly to XGBoost. A) Receiver-operator characteristic (ROC) curve for predicting CMap instances whose drugs were associated with autoimmunity. AUC: Area under the curve. B) ROC curve for predicting CMap instances whose drugs were associated with autoimmunity, considering only cell lines that performed well in validation. C) Per-drug scores for drugs associated with autoimmunity and control drugs. Scores are significantly different (p=0.009). D) Violin plots of per-drug scores as in (C) plus the per-drug scores of statins and ACEIs. Statins and ACEIs score significantly differently than control drugs (p=0.048) but not other autoimmune-inducing drugs (p=0.511). E) Boxplot displaying the prediction scores given to all FDA/EMA approved drugs by the model.FB) Violin plot of the prediction scores for drugs with at least two reports of SLE or CLE and for drugs not associated with lupus. The prediction score of drugs associated with lupus is significantly higher than other drugs (p = 0.010).

### The drug autoimmunity model is enriched with genes involved in cell cycling and proliferation

We extracted the genes used in the decision tree and ranked them by their overall order of importance in the model. We then used the top 500 genes to query the Molecular Signatures Database (https://www.gsea-msigdb.org/gsea/msigdb/; (Subramanian *et al*., 2005). We queried the following gene set collections, taking the most enriched gene set for each: Hallmark genes (H); chemical and genetic perturbations (CGP); BioCarta (CP:BIOCARTA); KEGG (CP:KEGG); PID (CP:PID); Reactome (CP:REACTOME); and WikiPathways (CP:WIKIPATHWAYS). We observed common patterns of enrichment, driven by 104 genes (52 genes, if excluding the chemical and genetic perturbations; Table S2). These pathways - cell cycle (3 collections), G2M checkpoint, minichromosome maintenance, E2F transcription factor network, and BRCA-correlated genes – are all linked by regulation of cell cycle, DNA replication, and proliferation.

## Discussion

In this paper, we have demonstrated that it is possible to use machine learning with an *in vitro* transcription perturbation dataset to predict drugs that can precipitate autoimmune conditions. Not only did our models accurately classify previously unseen perturbations (drug/cell line combinations), but they were also capable of predicting previously unseen drug classes. This is despite the strong likelihood that different mechanisms are responsible for different drugs causing autoimmunity, and/or that different autoimmune conditions can result from certain drugs. This is likely to be due to the ability of ensemble machine learning algorithms to capture complex feature relationships. Importantly, a similar approach could be easily employed in the preclinical phase of drug development. After initial capital investment, cost per sample for the L1000 platform, which is employed by CMap, is not much more than £1 ($1.50) (Subramanian *et al*., 2017). Therefore, following an initial small molecule screen, several (up to hundreds) of potential small molecules could be further screened against dozens of cell lines for autoimmune potential at limited cost. Our models could then provide additional information to be considered alongside target specificity, binding affinity, potential bioavailability, etc. to choose a lead compound.

The factors that make idiosyncratic autoimmune drug reactions difficult to study also present challenges for machine learning approaches. Knowing the “ground truth” (whether a molecule genuinely precipitates autoimmunity or not) affects both the accuracy of model training and our ability to interpret the model’s performance when tested against drug classes to which it was previously naïve. Even after we extracted drugs associated with CLE and SLE, the highest scoring of the novel drugs was sertraline, with a score of 0.896, which would easily classify it as being capable of precipitating autoimmunity. On the one hand, there are at least two case reports of lupus associated with sertraline (Hussain and Zakaria, 2008; Hill and Hepburn, 1998), suggesting that the model may be accurately classifying the drug. On the other hand, case reports are not definitive, and there is as yet insufficient information for lupus to be associated with sertraline in the various registries.

The biology revealed by examining the constituent genes of the model appears to centre on regulation of the cell cycle, DNA replication, and proliferation. It is important to understand that these are the genes used by the algorithm to make a decision, but it is not immediately clear whether there is overall activation of these genes, inhibition of these genes, or a mix of activation of some and inhibition of others. Nor is it trivial to infer this information from the model, which to some extent exists as a “black box”. For example, it is even possible that one gene may give a pro-autoimmune score when its expression is high in one tree, but low in another; context is important. More work will be required to better gauge the gene expression changes induced by drugs associated with autoimmunity and how they relate to the cell cycle. However, it is tempting to speculate that, because these pathways are necessary for lymphocyte activation and proliferation, the drugs associated with autoimmunity could alter the threshold of lymphocyte activation or could lead to an aberrant response to antigen presentation, including self-antigen. Better understanding the exact nature of the gene expression changes will lead to more refined hypotheses and experiments that can be used to confirm or refute these ideas.

There is more that could be done to refine our approach and improve the accuracy of its prediction. For example, as noted previously, other groups have had success with using structural data to predict DIA (Wu *et al*., 2021; Guo *et al*., 2022). It may therefore be possible to combine transcriptional with structural data. A better understanding of the underlying biology could help choose the most informative cell lines – which may not be as well-represented in the CMap database – and more accurate re-training of the model. Combining gene expression machine learning with case-control data from national patient registries, (such as the recent approach to identify drugs associated with lupus (Haugaard *et al*., 2020) may also improve accuracy, or better quantify uncertainty. Nevertheless, we believe that our approach could be incorporated into the drug discovery pipeline to help with selection of candidate molecules less likely to cause DIA.

## Acknowledgements

This work has been funded by the Medical Research Council Toxicology Unit (U.PA.PAPU.ANBK.FAAE.0000), and the Kay Kendall Leukaemia Fund Grant (KKL 1442). We acknowledge BioRender which was used to generate some of the figures presented in this publication.

