## Supplementary Information for "A machine learning approach to predict drug-induced autoimmunity using transcriptional data"

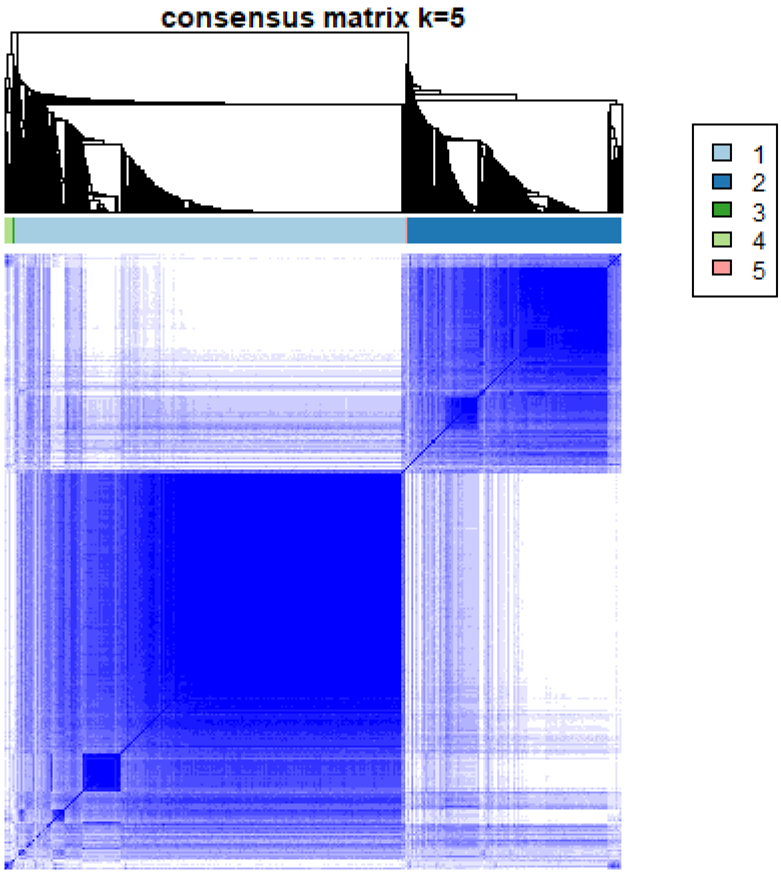


**Figure S1 Consensus clustering results when we specify five clusters.**

In comparison to the highly stable two-cluster solution shown in Fig 2A, specifying five clusters does not improve cluster stability when using consensus clustering, and clusters consist of very few CMap instances.
